# Advanced Cardiovascular Toxicity Screening: Integrating Human iPSC-Derived Cardiomyocytes with In Silico Models

**DOI:** 10.1101/2024.06.18.598996

**Authors:** Andrey Berezhnoy, Anastasiya Sinitsyna, Ivan Semidetnov, Vadim Naumov, Tatyana Sergeeva, Sergey Bakumenko, Mikhail Slotvitsky, Valeriya Tsvelaya, Konstantin Agladze

## Abstract

The pharmaceutical industry is evolving with the use of hiPSC-derived cardiomyocytes (hiPSC-CM) for in vitro cardiac safety screening. Traditional reliance on QT interval prolongation as a main cardiotoxicity marker is being challenged. In addition, Comprehensive In Vitro Proarrhythmia Assay (CiPA) initiative recommends using computer modeling and in silico platforms as more comprehensive approach for cardiotoxicity testing in conjunction with hiPSC-CM in vitro screening. Our study presents such an innovative platform that integrates in vitro hiPSC-CM propagation test with in silico models to assess cardiotoxicity. Utilizing the electrophysiological and morphological characteristics of hiPSC-CM, we offer a thorough evaluation of potential drug-induced cardiac risks by computer modelling. We show, using the example of lidocaine and other antiarrhythmics, that using a integrative experimental and computer platform, the possibility to correctly display the clinical manifestations of side effects in advance.

## 1. Introduction

To date, one of the most important markers of cardiotoxicity is the so-called QT toxicity. The correlation found between polymorphic ventricular tachycardias and action potential prolongation [1] has led in particular to the European CPMP Committee recommending routine QT interval prolongation testing for all pharmacological products [2]. On the other hand, it has become clear in recent years that action potential prolongation is not a reliable marker of potential arrhythmias and sudden ventricular death [3,4].

The most common and standardized test today is the hERG channel test (encoded by KCNH2) [5,6], but despite the mandatory requirement of this test, post-marketing drugs demonstrate the presence of cardiotoxicity. A systematic review showed that primary cardiotoxicity accounted for 74% of post-marketing drug withdrawals, such as arrhythmias (35%), cardiac damage (cardiomyopathy) (38%) and cardiovascular events (22%). In addition, during the preclinical to marketing and post-registration stages, cardiovascular problems accounted for 22% of treatment discontinuations [7]. To address this problem, the Comprehensive in vitro proarrhythmia analysis (CiPA) initiative was established [8]. In addition, the Japanese National Institute of Medical Sciences is also working on developing a more accurate method for clinical proarrhythmia risk prediction called Japan iPS Cardiac Safety Assessment JiCSA [9]. The study showed that the methods proposed in the initiatives are more accurate than current recommendations. The CiPA and JiCSA initiatives have demonstrated that cardiomyocytes induced from iPSCs (hiPSC-CM) can assess proarrhythmia risk. In addition, many reports show that hiPSC-CM assessments can be useful for assessing drug-induced toxicity and managing the development of new drugs in the preclinical phase. Long-term toxicity is a serious clinical problem because it can be unpredictable. Researchers have used hiPSC-CM to study the cardiotoxic effects of both existing and new drugs, including BCAs, anthracycline-doxorubicin, and low molecular weight kinase inhibitors [10,11].

Despite the necessity of the above-mentioned set of measures, a number of shortcomings of this initiative can be pointed out. First, it is the complexity of the whole complex of studies. Secondly, the study of drug action on ion channels can be performed only on isolated myocytes, and the density of delayed rectifier potassium currents, the traditional “culprits” of ventricular arrhythmias, is low in them [12]. Previously, to study the question of cardiotoxicity of drugs, we proposed a test representing an experimental iPSC model [13]. An isotropic monolayer of iPSC-derived cardiomyocytes was used as the excitable medium, stimulated by a punctate electrode that creates a spherical wavefront in the sample. A standard geometric obstacle (a straight section of the resulting cellular tissue perpendicular to the direction of wavefront propagation) with a large radius of curvature was present in the test as an initiator of wave breaking. The use of geometric obstacles is a classic wave breaking initiator that is widely used in the study of excitable media of various types [14]. Such a test can be expensive and often misses some of the mechanisms that lead to the occurrence of reentry at a standard obstacle.

We propose a new set of cardiotoxicity tests that combines a small number of in vitro experiments on hiPSC-CM with predictive modeling of the effects of substances in silico. We have shown that our model can qualitatively replace some of the expensive experiments for predicting cardiotoxicity and anti arrhythmogenicity of compounds on iPSCs when we include indicators from an experimental test of a compound. We also compared our final model with simpler electrophysiological models that do not take into account intercellular contacts and concluded that, with some limitations, such models can also replace part of the experimental testing. A new set of combined in vitro and in silico experiments was tested with potentially arrhythmogenic substances: lidocaine and endoxan. With the help of this complex it will be possible to finally solve the problem of directly linking the set of ionic currents or the shape of the action potential to the state of reentrant arrhythmias.

## 2. Materials and Methods

### 2.1. iPS cell lines and directed differentiation of iPSCs into cardiomyocytes

In this article we used 2 cell lines reprogrammed from a healthy donor that was described and fully characterized previously: iSMA6L and m34sk3 [13, 15]. Direct cardiac differentiation of iSMA6L was performed according to the original Gi-Wi protocol [16]. For m34sk3 the direct cardiac differentiation was modified according to protocol with 48h CHIR99021 incubating [17]. The first cell contractions were observed from day 9 of differentiation. Optical mapping occurred when the culture reached 50 days.

### 2.2. Drug stocks preparation

Lidocaine was used like an example of class I antiarrhythmic drugs as well as sodium channel blocker in 1 mM stock concentration. Drug dilutions in Tyrode’s solution were equilibrated at 37°C under 5% CO2 for 30 minutes before application to the wells.

The solution of cyclophosphamide (CP, Baxter) was used in this work as an example of a potentially arrhythmogenic drug. For CP dilutions 1 mM stock solution was added in Tyrode and preincubated the same way as lidocaine. We used 2 main concentrations of CP: 639 μM, 852 μM.

### 2.3. Samples preparation for arrhythmogenicity test

All samples were grown on Matrigel substrate in 24-well plates in the process of differentiation. Immediately before optical mapping, an acute unexcited obstacle was created on the culture to set the desired radius of curvature of the excitation wave front. The minimum size of the obstacle was limited by the thickness of the needle (20 μm). The methodology of sample creation is described in more detail in the previously published and tested arrhythmogenicity test [13].

### 2.4. Optical mapping part of arrhythmogenicity test

For optical mapping, we used a setup of an Andor iXon-3 high-speed light-sensitive camera (USA) mounted on an Olympus MVX-10 Macro-View optical stand (Olympus, Japan) with a corresponding Olympus fluorescence cube (similar to the cube for GFP -Olympus U-M49002XL). Fluo-4, AM dye (Invitrogen, USA) was used as a fluorescent dye to visualize the excitation wave conduction. Before optical mapping, the culture was incubated at 37°C for 30 minutes, and then the medium was replaced with Tyrode’s solution, which was brought to 37°C with stable pH. Optical mapping was performed in this Tyrode’s solution, maintaining the temperature condition for the cells at all times.

Two electrodes were used for tissue stimulation: a looped platinum wire reference electrode and a platinum wire stimulation point electrode. The stimulating electrode was placed ∼0.3 mm above the sample near its edge to minimize the spot of electrotonic excitation. The video was recorded at a speed of 130 frames per second and a spatial resolution of ∼0.1 mm/pixel. The pulses were in the form of a step with an amplitude of 4-6 mV, a duration of 20 ms, and a frequency of 1 to 4 Hz.

### 2.5. Immunofluorescent analysis and confocal microscopy

Immunofluorescent analysis was performed both on the layers of cells obtained during differentiation and on single cardiomyocytes to parameterize the Potts model. Single hiPSC-CMs samples were obtained by seeding differentiating samples with TrypLE solution (https://www.thermofisher.com/order/catalog/product/12605036) on Matrigel-coated glasses at a concentration of 50 thousand cells per ml. Cells were treated after 24h with Mitomycin-C to control cell division. After 7 days, the cells were fixed in 4% paraformaldehyde, permeabilized for 10 min in 0.4% Triton-X100 according to the protocol [18]. Cells were further incubated for 30 min in blocking buffer (1% bovine serum albumin in phosphate-buffered saline, PBS), overnight at 4°C with primary antibodies and for 1 h at room temperature with secondary antibodies. Cells were stained with antibodies for α-actinin, F-actin, Cx-43, and DAPI. Individual cells were imaged using an Olympus IX-71 fluorescence microscope and a Zeiss LSM 710 confocal scanning laser microscope. More complete immunofluorescent analysis and staining protocols are presented in previous works [13].

### 2.6. Computer modeling part of arrhythmogenicity test

#### 2.6.1. Electrophysiological model

The Kernik model was used for the experiments [19], adapted independently to derived induced cardiomyocytes for the myokit environment [20]. To mimic tissue heterogeneity, 10% of randomly selected elements had conductivities significantly lower than the standard conductivity (0.8 mS/μF) and were selected as 0.08 mS/μF. The size of an element was 50×50 μm and the time step of the simulations was 0.01 ms. The model used the Roush-Larsen difference scheme.

Simulations were performed on simulated virtual square cardiomyocyte specimens of 200×200 p×l (1×1 cm). The samples reproduced an obstruction in the form of a low conductance segment from the middle of one side of the square to its center, 2 pxl (100 μm) thick: 0.001 mS/μF for the obstruction region compared to 0.8 mS/μF for a normal element. A 25 pxl (1.75 mm) diameter circular stimulus was produced at the base of the obstacle (closer to the edge of the square). The duration of simulations was 7.2 seconds. In this case, the wave mode was investigated for the appearance of spiral waves on the obstacle, and the range of stimulation frequencies in which a stable reentry appeared was determined, i.e., the reentry corridor.

The values of sodium current blocking in this case were derived from experimental conduction velocities on monolayers. Details of the fitting and correlation of experimental and computer modeling are spelled out in the results, which represents the final test system. The corresponding potassium current blockages were obtained from the work [21], which were compared with the experimental results.

#### 2.5.2. Adapted iPSC Potts computer model

The Potts model, whose Hamiltonian is formulated as follows, was used to generate the samples:

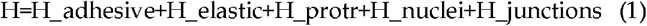

The formulation of the last term and the names of the constants and parameters are taken from [22, 23]. The processing of experimental data was carried out using ImageJ software. Basic metrics, used in the work, were obtained from confocal microscopy of single hiPSC-CMs:

- Cell area (area)
- The ratio of the cell area to the area of the circumscribed polygon
- Number of vertices of the circumscribed polygon
- Cell elongation

All metrics depend on all parameters in different ways, so the algorithm was developed for their joint search. For this work, the developed algorithm selected the optimization of the following parameters for different days of cell differentiation: Pdetach, GN, λ, LMAX, Nprotr, Vt, Jcell-cell, Jcell-MD, JCM-FB.

To compare the accuracy of parameter selection, the Jensen-Shannon divergence was used, which compared the distribution of experimental parameters with computer ones. In the calculation of the Jensen-Shannon divergence, the Kullback-Leibler distance was used. The processing was done using the Scikit-image and Feret libraries. The idea of the developed algorithm was based on Bayesian optimization and was to select parameters using machine learning (ML) with maximization of the accuracy of the resulting distribution function for each parameter. A detailed process of parameters selection is shown in Figure 1.

**Figure 1.**
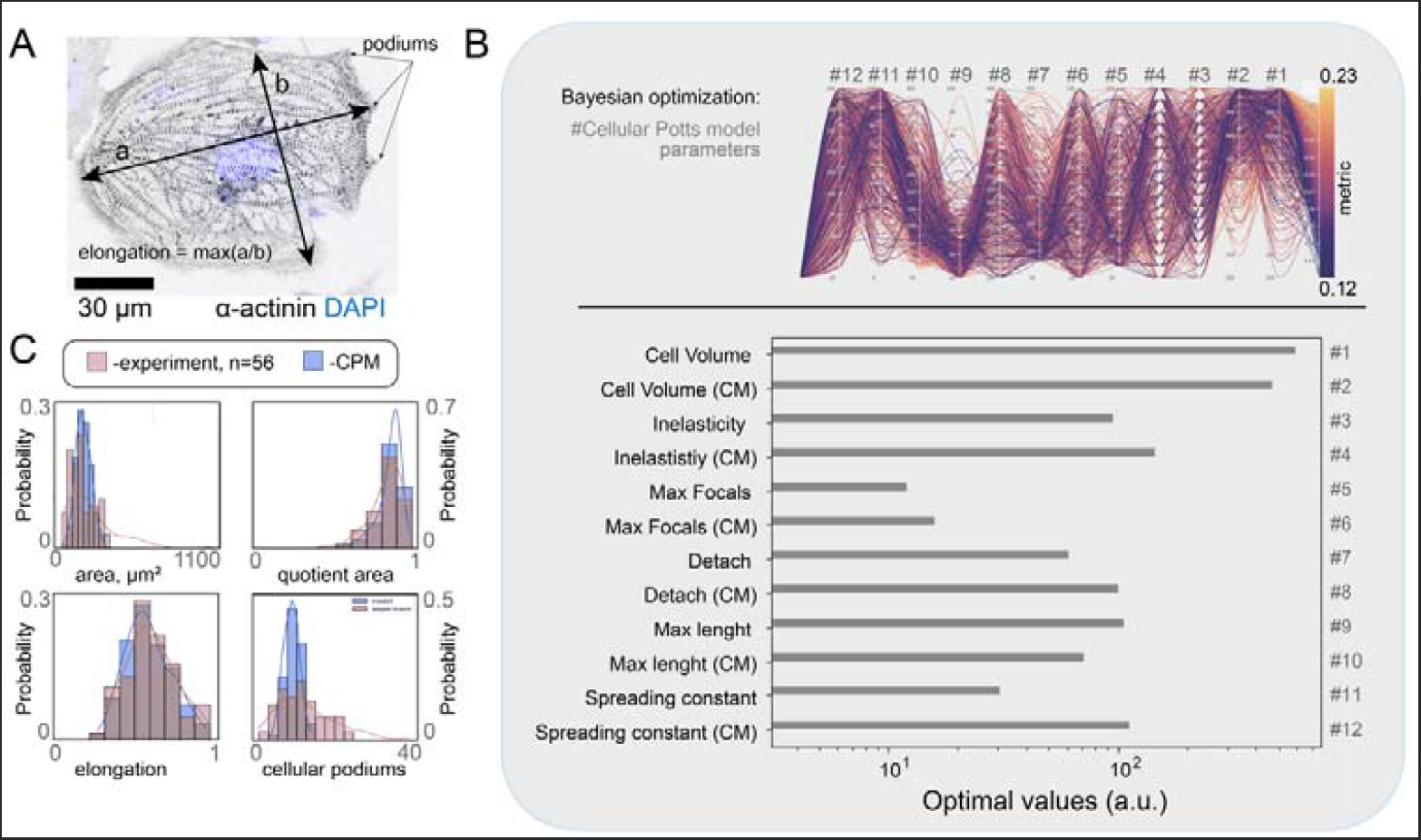
Example of compilation of Potts morphological model for hiPSC-CM by Bayesian optimization. A: Confocal image of single hiPS-CMs with 2 parameters of maximal length and width parameters of cell sizes a,b; B: Parameters selection using Bayesian optimization and their convergence with results of selection of morphology parameters with numbers from 1 to 12 for virtual cardiomyocytes in the Potts model; C: final convergence and selection of several parameters of the Potts model in comparison with experimental values.

The series of experiments carried out to study electrophysiology in tissue monolayers with cellular structure include 2 types of modeling: respectively, the Potts model to create the structure of the cardiac tissue and the model of cellular electrophysiology described in section 2.6.1. Tissue heterogeneity was achieved by introducing regions of different conductivities corresponding to cardiomyocytes, connexins and intercellular boundaries. The conductivities of these regions are correlated as 15/500/1000. The dimensions of the structural element of the monolayer are 5×5 μm.

Experiments to investigate the occurrence of reentry were performed on samples of square shape with a side length of 4 mm. The sample also reproduced an obstacle in the form of a 2 mm long segment, the conductivities of the elements of which are 0.001 of the corresponding conductivities of the normal regions. The location of the obstacle is similar to the case of the non-cellular model.

The simulation was performed in the open environment for cardiac modeling of electrophysiology in silico openCARP. The time step in the simulation calculations was 40 μs. The stimulation duration was 3 seconds. Stimulation was performed with a circular electrode at a point slightly below the base of the obstacle, radius 100 μm, stimulus amplitude 800 mV.

In experiments to determine the stimulation frequencies under CP exposure at which the excitation wave is assimilated by the virtual culture, simulations were performed on a strip of cells with dimensions of 0.5×2 mm without additional obstacles. Simulations of the effect of lidocaine on the tissue were performed similarly to the non-cellular only electrophysiological model. The effect of endoxan as a cell dissociator was simulated by decreasing the conductance of the intercellular junction by a factor of 5000 compared to the initial value. At the same time, the values of currents and conductances of other regions remained the same.

The calculations performed to simulate the tissue morphology were carried out on an IntelCore i7 CPU. Electrophysical calculations were made using Nvidia video cards of the Tesla series. Wolfram Mathematica was used to combine the morphological and electrophysiological models. The full version of the Potts model for cells differentiating into cardiomyocytes is available at the link: https://github.com/kalnin-a-i/pyVCT/tree/master. The plugin for optimizing parameters can be found at the link: https://github.com/violinist2802/Potts-optimization.

### 2.6. Data processing

All videos from optical mapping were processed in the ImageJ program. The activation and amplitude maps were built using the Wolfram Mathematica 9 program and Image J. The image processing from the confocal microscope was performed in the Zen program corresponding to the microscope software.

## 3. Results

### 3.1. Experimental arrhythmogenicity test

Not all arrhythmia mechanisms are reproducible on a single cell, so we previously presented an arrhythmogenicity test for different substances [13]. In the first experimental part of this paper, we reproduced this test to determine some parameters to add to the computer modeling, namely the conduction velocity. Our idea was also to do all the other experiments to compare them and the predicted arrhythmia mechanisms in the models with a minimal input of experimental data. In all our tests, we essentially evaluated the effect of stimulation frequencies on the conduction of the excitation wave. This resulted in 4 possible cases obtained by visualization of the test data: normal conduction without obstacle, normal conduction at a nonconducting obstacle, the precritical case in which the conduction wave detaches from the obstacle without reentry, and the critical case in which the conduction wave detaches from the obstacle with reentry (Figure 2).

**Figure 2:**
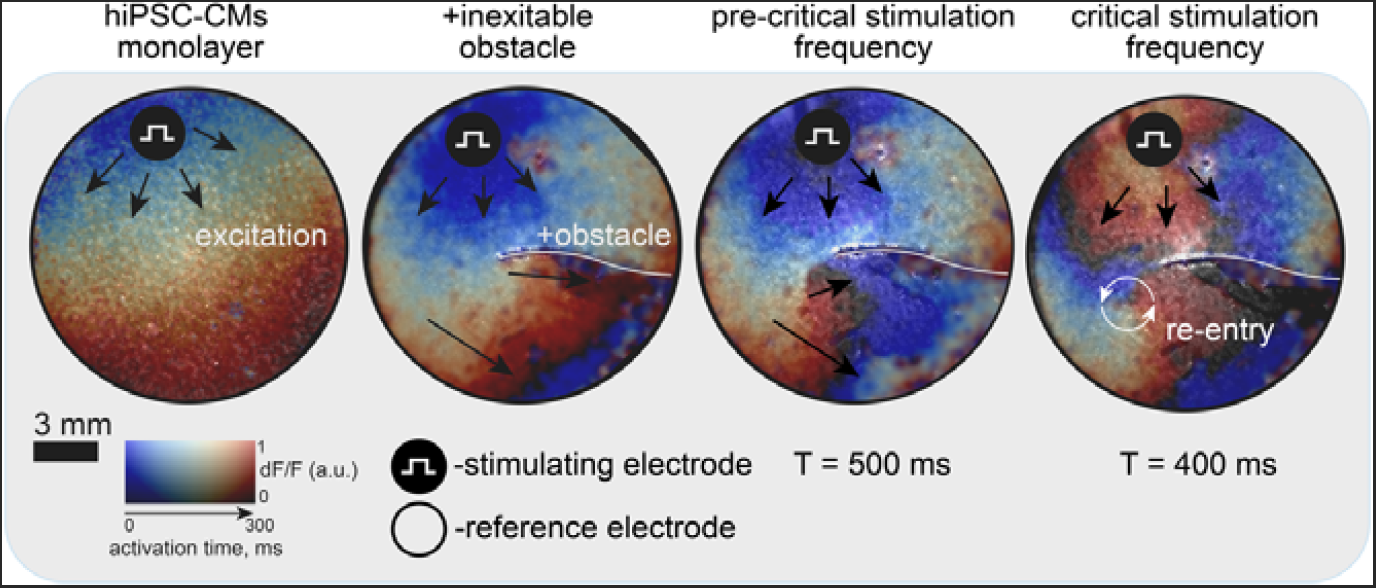
Schematic of the obstacle experiment. Four cases obtained in the experiment are shown: control of the excitation wave conduction without obstacle, normal conduction on a non-conducting obstacle; the pre-critical case, in which the excitation wave detachment from the obstacle is visible without the formation of a reentry; the critical case of the excitation wave detachment from the obstacle with the formation of a reentry.

Regarding the mechanism of reentry at the critical frequency, we considered 2 cases: direct wave detachment from an acute non-conducting obstacle with the occurrence of reentry; and spontaneous depolarizations during the repolarization phase of an action potential in cardiac myocytes or early afterdepolarizations (EADs). EADs are mostly not detectable in in vitro assays due to culture heterogeneity, but can be observed in vivo. In our case, we did not observe EADs as a cause of reentry in the experiment, but these cases appeared in silico, which completes the picture of arrhythmogenicity. In general, to validate the model and to understand whether an in silico test can reproduce experiments, we studied the effect of lidocaine. Its effect is known to be ambiguous. From the experiment, we measured the conduction velocity of the excitation waves under lidocaine. In the control, the velocity was about 5 cm/s with an error of 0.3 cm/s. At a lidocaine concentration of 50 μM, the velocity decreased and was 3.9 cm/s with an error of 0.28 cm/s. At a lidocaine concentration of 100 μM, the velocity decreased and was 3.3 cm/s with an error of 0.2 cm/s. At a lidocaine concentration of 200 μM, the velocity decreased and was 2.8 cm/s with an error of 0.15 cm/s. The other parameters of the in silico model were adjusted to these velocities at 1 Hz stimulation without obstruction and the arrhythmogenicity of lidocaine was determined. For comparison with the modeling results, the results were also obtained for the conduction of excitation waves on a non-conducting obstacle. We similarly measured CV rates for different CP concentrations. All experimental test data, with which computer simulation data were subsequently compared, are presented in the repository at the following link: 10.5281/zenodo.11441922

### 3.2. In silico electrophysiological modeling like a part of arrhythmogenicity test

The velocity depends on the currents and the intercellular contacts. Therefore, we identified ionic currents for the electrophysiological model from the velocities obtained from the experiment. We also looked at the blockade data and adjusted the currents for the desired velocities in the model so that the desired blockade currents matched the literature data on the patch-clamp assay. The simulation of drug effects on tissue was done by selectively blocking sodium and rapid potassium ion currents accordingly. Potassium blocks were taken from literature data, sodium was selected according to potassium and rate at a given drug concentration [21]. The final values of block for velocity adaptation when exposed to different concentrations of lidocaine are shown in Table 1.

**Table 1.** Similar blockade of sodium and rapid potassium ionic currents at different concentrations of lidocaine in the study model.

| Concentration x (mM) | Degree of ion channel blockade (GNa) | Degree of ion channel blockade (GKr) |
| --- | --- | --- |
| 0 | 0 | 0 |
| 50 | 0,53 | 0,14 |
| 100 | 0,68 | 0,25 |
| 200 | 0,81 | 0,4 |
| 300 | 0,86 | 0,5 |

In this way, we obtained an electrophysiological model for the conduction of excitatory waves across the modeled monolayer without taking cell morphology into account. The required velocities were obtained exclusively by correct selection of the variables for the ionic currents, so that in the case of lidocaine we do not need to take into account the change in intercellular contacts (Figure 3A). We also compared our obtained values with already published data where modeling on single cells showed the arrhythmogenicity of lidocaine. Our model showed good convergence with both our experiment and published experimental current data [21]. That is, changes in velocity are associated with changes in action potential (AP). We can observe in the simulation how the AP is prolonged by IKr and the role of calcium increases with increasing lidocaine concentration. Therefore, we were able to observe the effect of EAD and the increase in the probability of its occurrence even in a simplified electrophysiological model, in contrast to the experiment. That is, lidocaine is suitable to validate the relationship between currents and arrhythmogenicity in the simplified model (Figure 3B). We created a random distribution of diffusion coefficients and this was sufficient for modeling and predicting the experiment (Figure 3C).

**Figure 3.**
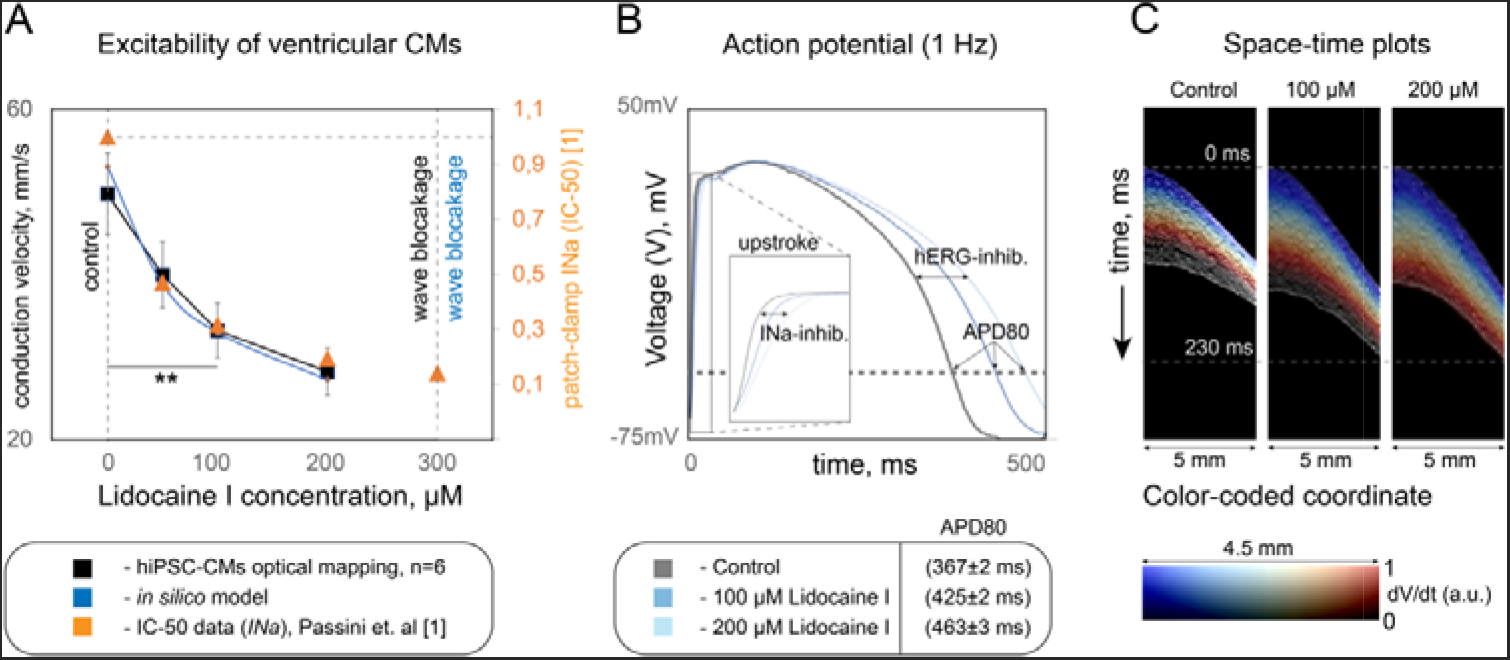
Excitation wave conduction by selected currents in an electrophysiologic model. A: Comparison of velocity conductance in the obtained simulation (blue), where the velocity was selected according to our experiment, in the experiment (black), in the simulation based on the published experiment (yellow), where the sodium conductance was set according to the patch-clamp data; B: Action potentials obtained in the modeling at different concentrations of lidocaine; C: Spatial and temporal sweeps of excitation wave conduction obtained in the model, from which one can see the inhomogeneities given by random diffusion and the difference in excitation wave conduction velocities

We then compared the performance of the resulting model in predicting a corridor of frequencies. We predicted this corridor of critical frequencies at which reentry occurs and saw that it was consistent with the experimental data. Without lidocaine, reentry occurred in the model in a corridor of stimulation duration from 350 ms to 374 ms, at a concentration of 100 μM from 397 ms to 424 ms, and at a concentration of 200 μM from 425 ms to 449 ms.

We observed that both reentry mechanisms were reproduced in the model: reentry at the barrier and reentry from early depolarization. Most importantly, the correlation between these two mechanisms is seen in this model. EADs do occur more frequently under lidocaine, as predicted on single cells, as shown in Figure 4. However, the contribution of EADs to the total number of reentries is very small compared to the probability of reentry at the barrier. It is also interesting to note that only at the barrier, lidocaine has an antiarrhythmic effect at 200 μm and a proarrhythmic effect at 100 μm. This is in full agreement with the results of our experiment. Thus, the test allows both to find the corridor of critical frequencies for the occurrence of reentry and to understand in advance the mechanism of reentry occurrence, for example in the presence of EAD.

**Figure 4.**
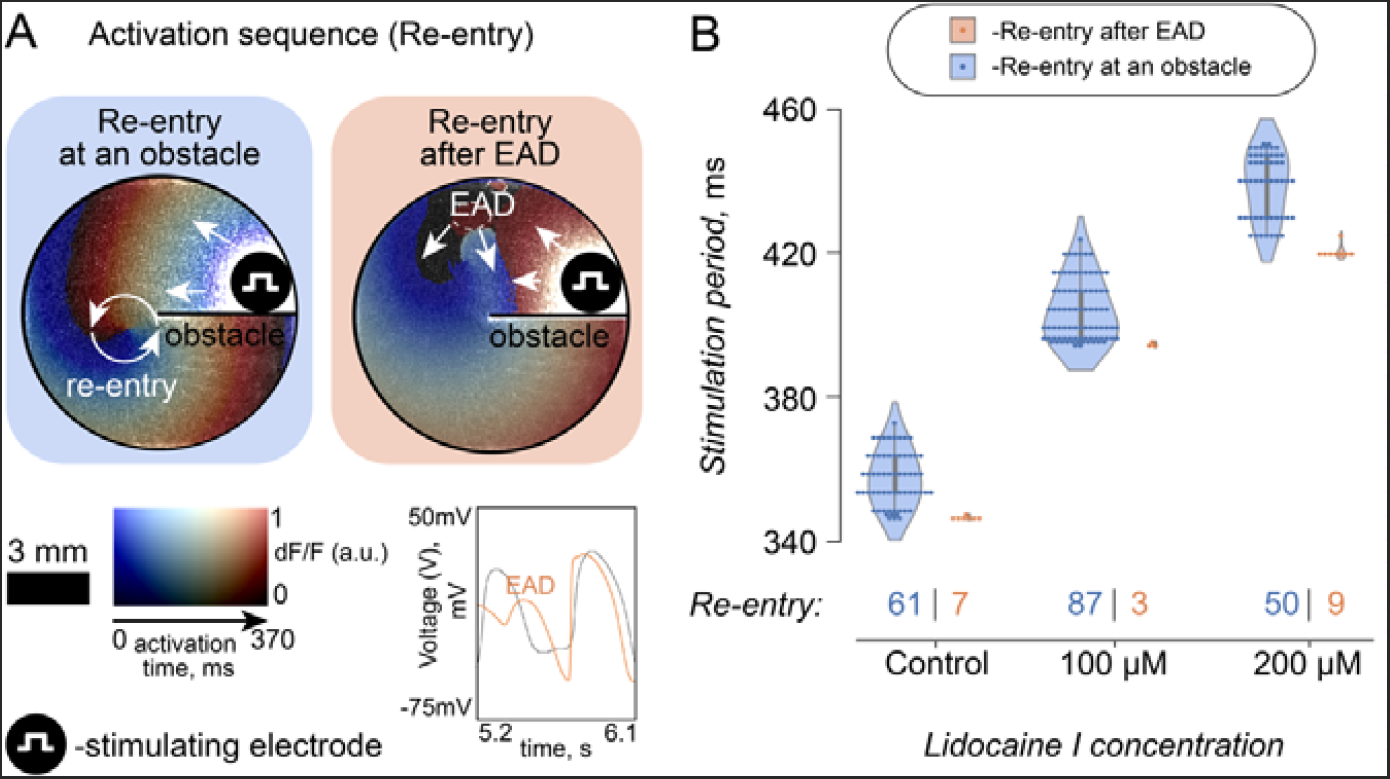
Results of predicting the occurrence of reentry in the obtained simple electrophysiologic model; A: Activation maps of possible 2 mechanisms of reentry occurrence obtained in the model: at the obstacle and in the case of EAD; B: The corridor of critical periods of stimulation (ms) at which reentries could form, with the amount of each mechanism’s contribution to their formation obtained in the model.

### 3.3 In silico Potts modeling like a part of arrhythmogenicity test

In the next step of the study, we complicated our electrophysiological model by adding parameters of tissue formation from hiPSC-CM using the Potts model. According to these methods, we measured area, elongation, described area and number of podia in isolated hiPSC-CMs of the m34sk3 lineage from confocal images of single hiPSC-CMs. Using Bayesian optimization, we were able to fit the mean and even the variance of the morphology parameter values for cellular modeling (Figure 1). Thus, we obtained a Potts model for adult hiPSC-CMs, which allowed us to move from a single-cell model to a 2D model.

Using the Potts model with the same electrophysiological model component, we also tested the corridor of frequencies at which reentry occurs and the possible mechanisms of this occurrence (Figure 5). We plotted a pie chart of the dependence of the probability of reentry occurrence at different stimulation frequencies on lidocaine concentration to compare these frequency corridors. To compare the corridors and to identify the y-axis as the coordinates of the stimulation frequencies, we made all period corridors uniform for control and lidocaine: we shifted the stimulation periods at which reentry occurred (T) by the MCR. For control, the MCR was 348 ms, for 100 μM lidocaine 395 ms, and for 200 μM 425 ms. We then divided the values of the stimulation periods, which represent the frequency corridors for the occurrence of reentry, by the length of the action potential. We were guided by the classical relationship that the corridor width is approximately 10% of the action potential length [24]. This relationship also holds for the frequency corridors we obtained in the Potts model, so the relationship holds for the Y-axis: 0<Y<0.1. The X-axis represents the number of spiral turns that fit into the 7.2 s computer simulation. The period step in the simulations was 1 ms.

**Figure 5.**
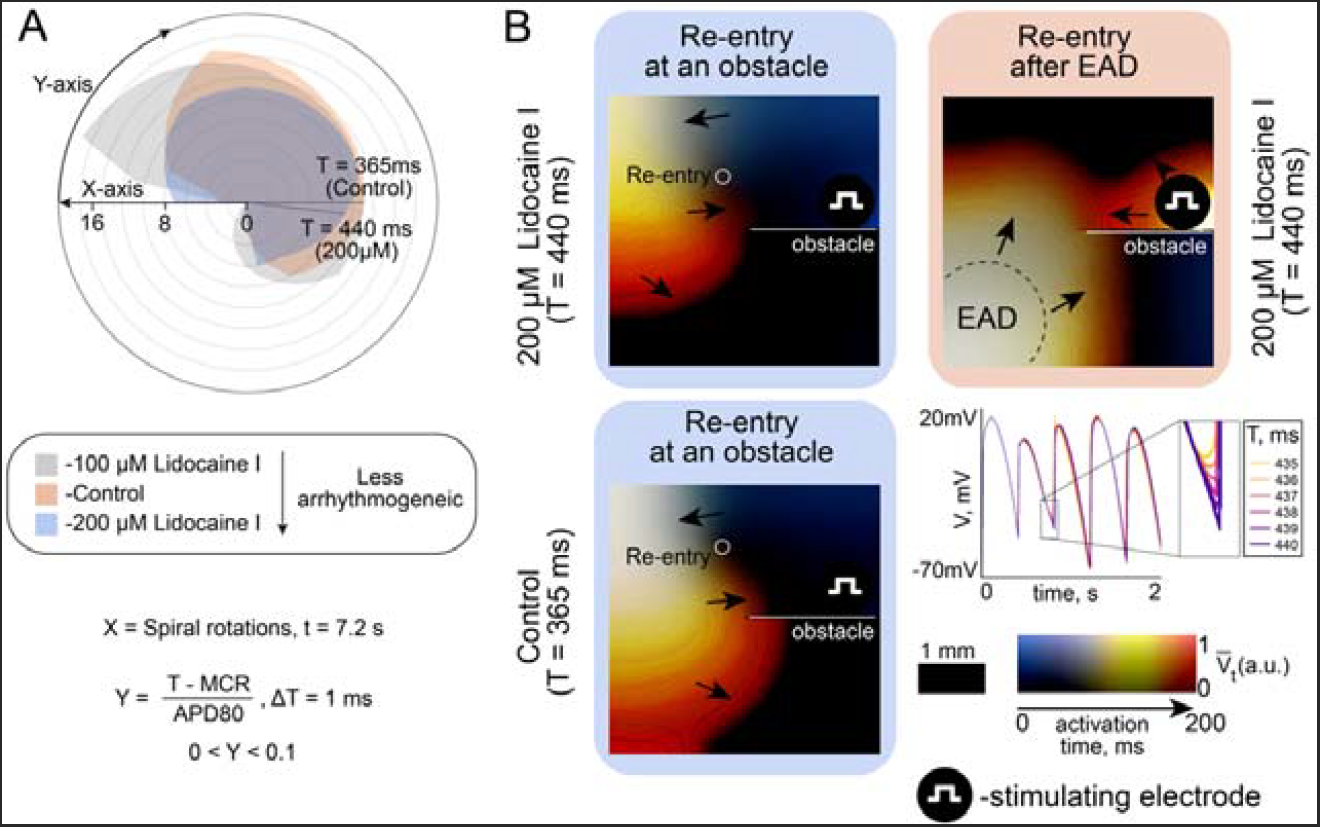
Frequency corridor modeling results for the occurrence of reentry at the obstacle and due to the effect of EAD occurrence. A: Pie chart comparing corridors of stimulation periods at which reentry occurs at different lidocaine concentrations and in the control, where X-number of spiral turns, for a fixed simulation length of 7. 2 ms, E-stimulation period, MCR-fixed shift value to bring the initial frequency corridor to a single origin, APD80-action potential duration measured at 80% repolarization; B: activation maps of excitation wave conduction across a hiPSC-CM monolayer modeled with the Potts model comparing stimulation periods on 200μM lidocaine from 435ms to 440 ms, i.e. in the resulting corridor of reentry occurrence. A comparison of the two reentry mechanisms on maps is presented.

Thus, we can clearly observe that at 200 μM lidocaine exhibits antiarrhythmic properties, although the probability of reentry remains. This is consistent with our experimental data, except that in the experiment we cannot observe the effect of EAD leading to reentry. Here, arrhythmogenicity was often accompanied by this effect, as we have shown in the activation maps in Figure 5B.

In the Potts model, we compared the model data obtained with the CP experiment, where the velocity decreased significantly under the influence of this substance (Figure 6). We analyzed the optical mapping data and took advantage of the fact that the Potts model is a complete simulation of layer formation and mapping. This allowed us to compare not only the velocity and frequencies, but also the inhomogeneity of the excitation wave conduction front, i.e., to suggest which structure is affected by the addition of CP and to model this effect. Therefore, we performed an experimental immunocytochemical study to investigate the changes in Cx43 under the influence of CP. We found no significant differences between control and CP incubated cells. Therefore, connexins and currents in the model were not altered by the action of CP. Thus, a new hypothesis was formed that all experimental differences in the conduction of the excitation wave under the influence of CP compared to the control are explained by cell dissociation. Our model, in contrast to a simple electrophysiological model, allowed us to test this putative mechanism of dissociation and to confirm this hypothesis. Qualitatively, in the Potts model, the velocity and the critical frequencies for the occurrence of reentry decreased, while the inhomogeneity of the front increased, similar to the experiment.

**Figure 6.**
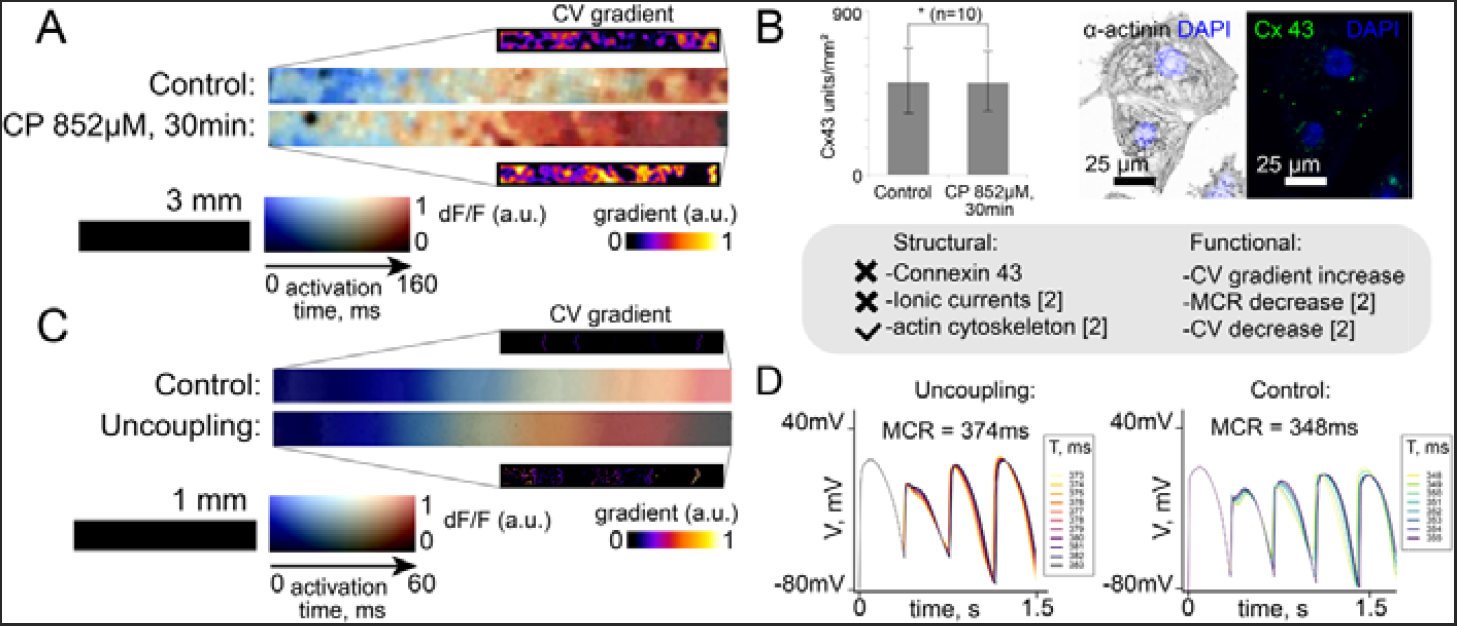
Computer simulation using the Potts model and comparison with CP-induced excitation wave conduction experiment. A: Data from an experiment with CP in a grown hiPSC-CM monolayer, activation maps of excitation wave conduction and its spatial gradient, where brightness indicates increased heterogeneities; B: Immunocytochemical analysis of cells after CP exposure, showing that there are no statistically significant differences in connexins in these cells compared to controls. At the bottom of panel B, hypotheses of rate-reducing mechanisms are presented, which are excluded or confirmed by modeling and experiment; C: Potts computer simulation with cell separation. The sweeps are plotted similarly to the experimental ones in panel A. It can be seen that the simulation qualitatively converges with the experiment in panel A; D: Convergence of the frequency corridor and its variation for control and at cell separation, simulating the effect of SR obtained with the Potts model.

## 4. Discussion

Despite the significant scientific background, the search for an optimal method to assess cardiotoxicity of pharmacological drugs is a real new task. The cardiotoxicity (arrhythmogenicity) test we present is not the only test for cardiotoxicity, but it has a number of advantages. The most common and standard test today is the hERG channel test (encoded by KCNH2) [5, 6], but despite the mandatory requirement of this test, drugs show the presence of cardiotoxicity after market release. A systematic review showed that primary cardiotoxicity accounted for 74% of post-marketing withdrawals, including arrhythmias (35%), cardiomyopathy (38%), and cardiovascular events (22%). In addition, cardiovascular problems accounted for 22% of discontinuations in the preclinical to marketing and post-approval phases [25]. An example of lidocaine and CP as potential antiarrhythmic and proarrhythmic agents has been reviewed in this paper [21].

In recent years, most of the work devoted to the practical application of mathematical models in computational cardiology has been conducted in two related areas: patient-specific modeling and drug cardiotoxicity assessment [26, 27]. In this work, we also proposed a common cardiotoxicity testing system and showed that even simple electrophysiological models can avoid some of the expensive experiments. Furthermore, we have shown that not all effects such as EAD can be detected in vitro. However, such effects are potentially arrhythmogenic when the drug is ingested. In our work, this is demonstrated in 2 models: a simplified electrophysiological model and a Potts cell model combined with the same electrophysiological model.

Remodeling at the cellular level has not been considered in models representing arrhythmogenicity testing. Although model personalization has been successfully applied in the study of atrial fibrosis at the cellular level [28, 29]. In our study, we complemented the electrophysiological model with morphological cell-specific modeling This allowed us to predict the mechanism of action of a compound at the Level of intercellular junctions and cell structures. This means that we can model, among other things, cellular dissociation that does not occur through gap junctions. We demonstrated this in our work using CP as an example. Our computer simulation data agreed with the repeated experiment presented in the paper and a previously published test of CP on hiPSC-CM monolayers [30].

In addition, we analyzed the effect on arrhythmogenicity of the action potential change obtained from the currents. As a result, we found that both models reproduced the experimental data. That is, the computer model can replace part of the experiments if we choose parameters specific to the experiments already performed. An interesting result of the work can be considered that we generally confirmed and reproduced the experiments from the arrhythmogenicity test, and thus were able to show both proarrhythmogenic and anti-arrhythmogenic side of lidocaine, in contrast to previously published works [21]. Our study demonstrated the importance of using 2D drug testing in hiPSC-CM.

## 5. Conclusions

We were able to make and validate on new and already published results a new joint in vitro and in silico test system, including a computerized cellular model of hiPSC-CM monolayer, which will allow unambiguous interpretation of pro- and anti-arrhythmogenic effects of substances with known effects on both membrane ion channels and non-intercellular structures. In the course of the study we have shown that the availability of such a test and its two-dimensionality is necessary for the correct evaluation of some known drugs - lidocaine. For example, on single cells it got into proarrhythmics, because only EADs could be evaluated on single cells, but not the totality of important interactions, including those with nonconducting obstacles. We have also shown that even simple electrophysiologic models effectively complement and replace part of in vitro experiments by predicting the same arrhythmogenicity effects. The test will allow high-throughput screening of any substance or combination of substances. To use it, it is sufficient to know a portion of the data from an electrophysiologic study such as a patch clamp. The test will predict a corridor of frequencies and the main mechanisms of arrhythmogenesis in this corridor, which can already be effectively tested with a minimum number of experiments on hiPSC-CM monolayers, since it imitates them directly.

## Author Contributions

Conceptualization, A.B., V.T., M.S., K.A., A.E., O.A., I.A., A.L.; methodology, A.B., A.S., I.S., V.N., T.S., M.S., V.T. ; software, A.B., A.S., I.S., V.N., T.S., M.S., V.T.; validation, A.B., A.S., I.S., V.N., T.S., M.S., V.T..; formal analysis, A.B., A.S., I.S., V.N., T.S., M.S., V.T.; investigation, A.B., A.S., I.S., V.N., S.B., T.S., M.S., V.T., K.A., A.E., O.A., I.A., A.L.; resources, A.B., A.S., I.S., V.N., S.B., T.S., M.S., V.T.; data curation, A.B., A.S., I.S., V.N., T.S., M.S., V.T..; writing—original draft preparation, A.B., A.S., M.S., V.T.; writing—review and editing, A.B., M.S., V.T., K.A., A.E., O.A., I.A., A.L.; visualization, A.B., A.S., M.S., V.T.; supervision, A.B., M.S.,V.T., K.A., A.E., O.A., I.A., A.L.; project administration, A.B., M.S., V.T., K.A., A.E., O.A., I.A., A.L.; funding acquisition, V.T., K.A., A.E., O.A., I.A., A.L.. All authors have read and agreed to the published version of the manuscript.

## Institutional Review Board Statement

The study was conducted in accordance with the Declaration of Helsinki, and approved by the Institutional Animal Care and Use Committee of M. F. Vladimirsky Moscow Regional Clinical Research Institute (protocol No. 4, 4 March 2021) and by the Moscow Institute of Physics and Technology Life Science Center Provisional Animal Care and Research Procedures Committee (Protocol No. A2-2012-09-02).

## Informed Consent Statement

Not applicable

## Data Availability Statement

The full version of the Potts model for cells differentiating into cardiomyocytes is available at the link: https://github.com/kalnin-a-i/pyVCT/tree/master. The plugin for optimizing parameters can be found at the link: https://github.com/violinist2802/Potts-optimization.

The electrophysiological model used in the work can be found at the link: https://github.com/CardioBioLab/iPSC_model

All simulations obtained in the work can be found on the disk: https://drive.google.com/drive/folders/1q4Rfg5iWcyZDvrhclH9pL3EbFTMm5ZFk

## Acknowledgments

The work was mainly supported by the Ministry of Science and Higher Education of the Russian Federation in the scope of the government assignment (Agreement 075-03-2023-106 13 January 2023). This research was funded by the Ministry of Science and Higher Education of the Russian Federation (Grant Number: 075-15-2022-310) and RUDN University Strategic Academic Leadership Program and M.F. Vladimirsky Moscow Regional Clinical Research Institute state grant #55. So, we would like to express special gratitude to the administration of the ITMO university and MIPT for the financial support of the authors. We would also like to express special gratitude to the administration of the «Tatneft» company for supporting the project.

## Disclaimer/Publisher’s Note

The statements, opinions and data contained in all publications are solely those of the individual author(s) and contributor(s) and not of MDPI and/or the editor(s). MDPI and/or the editor(s) disclaim responsibility for any injury to people or property resulting from any ideas, methods, instructions or products referred to in the content.

## References

1. Straus SM, Sturkenboom MC, Bleumink GS, et al. Non-cardiac QTc-prolonging drugs and the risk of sudden cardiac death. Eur Heart J 2005;26:2007–12

2. Committee for Proprietary Medicinal Products (CPMP/986/96) London: EMEA; Points to consider: The assessment of the potential for QT interval prolongation by non-cardiovascular medicinal products. December 17, 1997

3. Icilio Cavero & Henry Holzgrefe (2014) Comprehensive in vitro Proarrhythmia Assay, a novel in vitro/in silico paradigm to detect ventricular proarrhythmic liability: a visionary 21st century initiative, Expert Opinion on Drug Safety, 13:6, 745–758, DOI: 10.1517/14740338.2014.915311,

4. Meera Varshneya, Xueyan Mei, Eric A. Sobie. Prediction of arrhythmia susceptibility through mathematical modeling and machine learning. Proceedings of the National Academy of Sciences Sep 2021, 118 (37) e2104019118; DOI:10.1073/pnas.2104019118

5. Gintant, G.. An Evaluation of hERG Current Assay Performance: Translating Preclinical Safety Studies to Clinical QT Prolongation. Pharmacol. Ther. 2011, 129, 109–119.;

6. Pourrier, Marc, and David Fedida. The emergence of human induced pluripotent stem cell-derived cardiomyocytes (hiPSC-CMs) as a platform to model arrhythmogenic diseases. International journal of molecular sciences. 2020, 21.2, 657

7. Tani, Hidenori, and Shugo Tohyama. Human engineered heart tissue models for disease modeling and drug discov-ery.” Frontiers in Cell and Developmental Biology. 2022, 10, 855763

8. Yang, X., and Papoian, T.. Moving beyond the Comprehensive In Vitro Proarrhythmia Assay: Use of Human-Induced Pluripotent Stem Cell-Derived Cardiomyocytes to Assess Contractile Effects Associated with Drug-Induced Structural Cardiotoxicity. J. Appl. Toxicol, 2018, 38, 1166–1176

9. Kanda, Y., Yamazaki, D., Osada, T., Yoshinaga, T., and Sawada, K.. Development of Torsadogenic Risk Assessment Using Human Induced Pluripotent Stem Cell-Derived Cardiomyocytes: Japan iPS Cardiac Safety Assessment (JiCSA) Update. J. Pharmacol. Sci., 2018, 138, 233–239

10. Sakamoto, K., Sakatoku, K., Sugimoto, S., Iwasaki, N., Sano, Y., Yamaguchi, M., et al.. Continued Exposure of Anti-cancer Drugs to Human iPS Cell-Derived Cardiomyocytes Can Unmask Their Cardiotoxic Effects. J. Pharmacol. Sci, 2019, 140, 345–349.;

11. Gunawan, Felix, Rashmi Priya, and Didier YR Stainier. “Sculpting the heart: Cellular mechanisms shaping valves and trabeculae.” Current opinion in cell biology 73 (2021): 26–34.

12. Rajamani S, Anderson CL, Valdivia CR, Eckhardt LL, Foell JD, et al. (2006) Specific serine proteases selectively damage KCNH2 (hERG1) potassium channels and I(Kr). Am J Physiol Heart Circ Physiol 290: H1278–1288

13. Slotvitsky, Mihail, et al. “Arrhythmogenicity test based on a human-induced pluripotent stem cell (iPSC)-derived cardiomyocyte layer.” Toxicological Sciences 168.1 (2019): 70–77

14. Kléber A. G., Rudy Y. Basic mechanisms of cardiac impulse propagation and associated arrhythmias //Physiological reviews. – 2004. – T. 84. – No. 2. – C. 431–488

15. Slotvitsky, M. M., et al. “Formation of an electrical coupling between differentiating cardiomyocytes.” Scientific reports 10.1 (2020): 7774

16. Lian, Xiaojun, et al. “Directed cardiomyocyte differentiation from human pluripotent stem cells by modulating Wnt/β-catenin signaling under fully defined conditions.” Nature protocols 8.1 (2013): 162–175

17. Burridge, Paul W., et al. “Chemically defined generation of human cardiomyocytes.” Nature methods 11.8 (2014): 855–860

18. Podgurskaya, A. D. et al. The Use of iPSC-Derived Cardiomyocytes and Optical Mapping for Erythromycin Arrhythmogenicity Testing //Cardiovascular toxicology. 1–11 (2019)

19. Kernik, Divya C., et al. “A computational model of induced pluripotent stem-cell derived cardiomyocytes incorporating experimental variability from multiple data sources.” The Journal of physiology 597.17 (2019): 4533–4564

20. Clerx, M., Collins, P., De Lange, E., & Volders, P. G. (2016). Myokit: a simple interface to cardiac cellular electrophysiology. Progress in biophysics and molecular biology, 120(1-3), 100–114.

21. Passini, Elisa, et al. “Human in silico drug trials demonstrate higher accuracy than animal models in predicting clinical pro-arrhythmic cardiotoxicity.” Frontiers in physiology 8 (2017): 668.

22. Kudryashova, Nina, et al. “Self-organization of conducting pathways explains electrical wave propagation in cardiac tissues with high fraction of non-conducting cells.” PLoS computational biology 15.3 (2019): e1006597.

23. Kudryashova, Nina, et al. “Virtual cardiac monolayers for electrical wave propagation.” Scientific reports 7.1 (2017): 7887.

24. Cabo, Candido, et al. “Vortex shedding as a precursor of turbulent electrical activity in cardiac muscle”. Biophysical Journal 70.3 (1996): 1105–1111

25. Tani, Hidenori, and Shugo Tohyama. Human engineered heart tissue models for disease modeling and drug discov-ery.” Frontiers in Cell and Developmental Biology. 2022, 10, 855763

26. Trayanova, N. A. (2011). Whole-heart modeling: applications to cardiac electrophysiology and electromechanics. Circulation research, 108(1), 113–128;

27. Boyle PM, Zghaib T, Zahid S, et al. Computationally guided personalized targeted ablation of persistent atrial fibrillation. Nat Biomed Eng. 2019;3(11):870–879. doi:10.1038/s41551-019-0437-9

28. Syed Z, Vigmond E, Nattel S, Leon LJ. Atrial cell action potential parameter fitting using genetic algorithms. Med Biol Eng Comput. 2005;43: 561–571. pmid:16411628;

29. Bot CT, Kherlopian AR, Ortega FA, Christini DJ, Krogh-Madsen T. Rapid Genetic Algorithm Optimization of a Mouse Computational Model: Benefits for Anthropomorphization of Neonatal Mouse Cardiomyocytes. Front Physiol. 2012;3. pmid:23133423

30. Podgurskaya, A. D., et al. “Cyclophosphamide arrhythmogenicitytesting using human-induced pluripotent stem cell-derived cardiomyocytes.” Scientific reports 11.1 (2021): 2336.

